# The effects of arm swing amplitude and lower-limb asymmetry on gait stability

**DOI:** 10.1101/664565

**Authors:** Allen Hill, Julie Nantel

## Abstract

Changes to arm swing and gait symmetry are symptomatic of several pathological gaits associated with reduced stability. The purpose of this study was to examine the relative contributions of arm swing and gait symmetry towards gait stability. We theorized that actively increasing arm swing would increase gait stability, while asymmetric walking would decrease gait stability. Fifteen healthy, young adults (23.4 ± 2.8 yrs) walked on a split-belt treadmill under symmetric (1.2 m/s) and asymmetric walking (left/right, 5:4 speed ratio) with three different arm swings: held, normal, and active. Trunk local dynamic stability, inter-limb coordination, and spatiotemporal gait variability and symmetry were measured. Active arm swing resulted in improved local trunk stability, increased gait variability, and decreased inter-limb coordination (*p* < .013). The changes in local trunk stability and gait variability during active arm swing suggests that these metrics quantify fundamentally different aspects of stability and are not always comparable. Split-belt walking caused reduced local trunk stability, increased gait variability, and increased lower limb asymmetry (*p* < .003). However, the arm swing symmetry was unaffected by gait asymmetry, this suggests that deficits in gait stability in pathological gaits may be linked to increases in gait asymmetry rather than increases in arm swing asymmetry.

## Introduction

Arm swing during gait has been shown to have both passive and active components [1, 2]. While the exact interplay between these components in arm swing is still unknown, the small torques calculated at the shoulder indicate that passive, pendular like oscillations are likely dominant in the formation of normal arm swing. However, muscle activity at the shoulder is persistent, even during conditions of restricted arm movements (e.g. bound arms) [2]. This activity is thought to arise from central pattern generators important in the formation of normal gait patterns. Indeed, there appears to be a neurological link between the oscillatory movements of the arms and legs [3, 4]. The purpose of this neural connection, and more broadly of the ubiquitous presence of arm swing in healthy human gait, is presently unclear. Past research has linked arm swing during gait to decreased metabolic cost [5], decreased vertical ground reaction forces [6], and increased stability [7, 8].

Interestingly, Ortega et al. (2008) found no difference in the metabolic cost of gait between normal and held arm swing conditions when walking with external stabilization. Therefore, it was posited that the increased metabolic rate when walking without both arm swing and external stabilization was due to an increased effort in maintaining stability [5]. Further studies into this link between arm swing and gait stability have found conflicting results. Some studies have found decreases in stability when arm swing was prevented (either by binding the arms or holding them still) as compared to normal swing by analysing step width variability [9], harmonic ratios [10], and local dynamic stability [8]. Yet, others found no effect on stability when comparing bound and normal swing using local dynamic stability [7] and harmonic ratios [10]. Some have found increases in local dynamic stability when comparing active arm swing (e.g. increased swing amplitude) conditions to normal swing [11]. To our knowledge, no previous studies have compared the effects of normal, held and active arm swing conditions on gait stability within the same population.

A concept from dynamical systems theory, local dynamic stability uses an estimate of the maximal Lyapunov exponent (MLE), a measure of chaos, to quantify stability where a smaller MLE represents an increase in stability and a larger MLE represents a decrease in stability. Dingwell et al. were the first to use the MLE to quantify stability during steady-state gait and found it to be a more sensitive measure of stability than traditional gait variability measures when comparing overground walking to walking on treadmills in a population of healthy young adults [12]. Further studies have supported its use as a measure of stability; van Schooten et al. demonstrated the feasibility of MLE as an indicator of gait stability by using galvanic vestibular stimulation to reduce the stability of healthy young adults [13]. Looking at the local dynamic stability of lower limb joint angles [14] and of the trunk velocity [15], researchers also reported greater trunk stability with increased walking speed. Finally, older adults show significant decreases in gait stability compared to young adults when measured with the MLE [16].

Other metrics commonly used to measure gait stability are gait spatiotemporal variability metrics including the means and standard deviations of the length, width, and time of strides and steps [12, 17]. Increases in these gait variability measures are associated with increased fall risk in older adults as well as people with Parkinson’s disease (PD) [17–20].

Asymmetry in gait has also been linked to decreases in gait stability. Individuals with pathological gait, such as those with PD, often demonstrate lower limb asymmetry as well as changes in arm swing amplitude and symmetry when compared to age-matched controls [20]. Studies also link this asymmetric gait pattern to increased risk of falls [21]. In healthy young adults, asymmetric gait induced via split-belt treadmills is associated with decreased stability measured using margins of stability [22, 23]. However, the relationship between asymmetry in the upper and lower limbs and changes in stability is not well understood.

The purpose of this study was to further examine the role of arm swing and lower limb symmetry on gait stability in young healthy adults using common gait variability measures (mean and standard deviation of step length and width) and local dynamic stability. Based on previous literature, we hypothesized that active arm swing would lead to greater stability and that asymmetric walking would be less stable than symmetric walking.

## Materials and Methods

### Subjects

An *a priori* power analysis performed using SigmaPlot 12.5 (Systat Software, San Jose, CA) indicated that 12 participants would result in adequate statistical power when set at *β* = 0.8. To account for possible attrition, fifteen healthy, young adults (8 male, 23.4 ± 2.8 years (mean ± s.d.); 72.3 ± 13.5 kg; 170.2 ± 8.1 cm) from the Ottawa area were recruited. Subjects were excluded based on the presence of any recent (< 6 months) musculoskeletal injuries, or any chronic neurological or orthopaedic disorders that could affect gait. The study was carried out in compliance with the Tri-Council Policy statement; Ethical Conduct for Research Involving humans; The International Conference on Harmonization – Good Clinical practice: Consolidated Guideline; and the provisions of the Personal Health Information Protection Act 2004. The study was approved by the Ottawa Health Science Network Research Ethics Board (20170291-01H) as well as by the University of Ottawa Research Ethics Board (A06-17-03). All subjects gave written, informed consent prior to participation.

### Protocol

A 57 marker set was used to capture kinematic data [24] in the Computer Assisted Rehabilitation Environment (CAREN) (CAREN-Extended, Motekforce Link, Amsterdam, NL) at the Ottawa Hospital Rehabilitation Centre. The CAREN system combines a 6 degree of freedom platform with an integrated split-belt treadmill (Bertec Corp., Columbus, OH) and a 12 camera Vicon motion capture system (Vicon 2.6, Oxford, UK). Kinematic data was captured at 100Hz, and kinetic data was collected at 1000Hz. Participants were asked to walk on the split-belt treadmill at a speed of 1.2 m/s. Each trial had a duration of 200s and the first 25s of each trial were removed to allow the treadmill to reach the set speed and the participant to reach a steady-state. The three arm swing conditions were: held, normal, and active. For the held swing condition, participants were instructed to hold their arms at their sides in a relaxed manner, without swinging or stiffness. The instructions for the active swing condition were to swing their arms such that the arm was horizontal when each forward swing peaked. For the asymmetric walking condition, the right treadmill belt speed was set to 80% of the left side, or 0.96 m/s; the treadmill asymmetry ratio was chosen to mimic the asymmetries developed in older adults and other pathological populations [21, 25]. The combination of all arm swing and symmetry conditions were randomized for each participant; each condition was performed once.

### Analysis

All data was imported into Visual3D v6 (C-Motion, Germantown, MD) and filtered using a 4th order, zero-lag low-pass Butterworth filter with a cutoff frequency of 12Hz and 10Hz for the kinematic and kinetic data, respectively. Heel strikes were identified with a logistic classification model which used ground reaction forces from both force plates and kinematic data (feet position relative to the pelvis, feet velocity–relative to both the pelvis and the laboratory, feet acceleration, and both knee angles); all heel strikes were manually verified and corrected as necessary. Trunk linear and angular velocities, feet centre of mass positions, and left and right shoulder and hip angles in the sagittal plane were exported from Visual3D and further data analyses were done in Julia (v1.0.3) [26] using custom code. All measures were calculated using 125 strides of data.

Arm swing amplitude was calculated as the average sagittal range of motion of the shoulder angle. Continuous relative phase (CRP) between contralateral arm-leg pairs was calculated as recommended by Lamb and Stockl (2014) by first centering the amplitude of shoulder and hip angles, then calculating the phase angle for each signal using the signal and its Hilbert transform, shown in Eq (1–2). Finally, the CRP was calculated between the two signals as shown in Eq (3) [27]. We calculated the average of the ensemble standard deviation of CRP, hereafter referred to as MSDCRP, to quantify coordination as the interstride variability of CRP [27, 28].

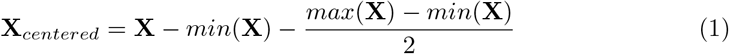

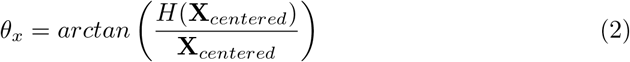

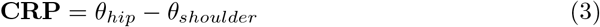

Note that the extrema, *min* and *max*, in Eq (1) were the average extrema of the 125 strides analysed. In Eq (2), *H*(**X**) represents the imaginary part of the analytic signal produced by the Hilbert transform. Circular means and standard deviations were used to reduce the CRP data to MSDCRP [29].

Average step width and step width variability were calculated as the average and standard deviation of the mediolateral distance between successive heel strikes. Average step length and step length variability were calculated as the average and standard deviation of the anteroposterior distance between successive heel strikes [17]. Step length metrics were analysed separately for each foot. This was necessary to avoid confounding effects in the asymmetric gait condition, where the left and right steps will naturally have 2 distinct lengths. Left steps are defined as the left heel strike following a right leg stance; right steps are defined oppositely.

Arm swing asymmetry, and step asymmetry-both temporal and spatial-were calculated as

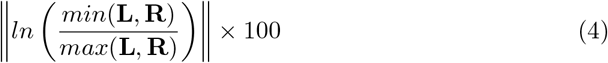

where **L** and **R** are the metrics for the left and right sides, and where a result of zero represents perfect symmetry [25]. Temporal step asymmetry was calculated using step time, and spatial step asymmetry was calculated step length.

### Local dynamic stability

Local dynamic stability, measured as the maximum finite time Lyapunov exponent (MLE), characterizes the average logarithmic divergence of two trajectories from infinitesimally close initial conditions. The MLE was calculated with Rosenstein’s method [30] using 125 strides of data interpolated to a length of 12,500 samples for an average stride length of roughly 100 points [31]. Note that 125 strides were analysed instead of the more common number of 140 strides because an unacceptable number of trials in the active swing condition did not contain an adequate number of strides. Data was reconstructed in a 12D state space of the form

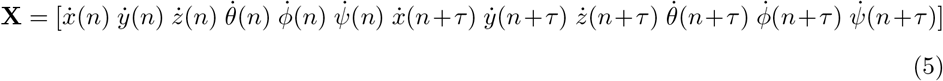

using the linear and angular velocity of the trunk and their 25 samples delayed signals [32]. To account for the differing units, the linear and angular velocities were normalized to unit variance [33]. Only the “short term” maximum finite time Lyapunov exponent, *λ*_*S*_, was used, which is defined as the slope of the divergence curve from 0 to 0.5 strides, inclusive [31, 32].

### Statistical Analysis

A two-way (Arms×Symmetry) repeated-measures (RM) ANOVA was performed on the following variables: left and right arm swing amplitude, arm swing asymmetry, average right and left step length and width, spatial and temporal step asymmetry, right and left step length and width variability, and MLE. One outlier was identified and removed for left step length variability and temporal step asymmetry. A two-way (Arms×Symmetry, normal and active swing vs. both symmetry conditions) RM ANOVA was performed on MSDCRP for both arm-leg pairs. Significance level was set *a priori* at *α* = 0.05. A Bonferroni correction was used for all post-hoc tests performed. Assumption of normality was confirmed using a Kolmogorov-Smirnov test and a Greenhouse-Geisser *p* was reported when Mauchly’s Test of Sphericity was violated.

## Results

### Arm swing range of motion and coordination

Table 1 contains the results of the arm swing range of motion (RoM) and MSDCRP for both contralateral limb pairs. A significant effect of arm swing was found for the left (*F* (2, 28) = 387.36, *p* < .001, 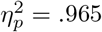), and right (*F* (2, 28) = 362.65, *p* < .001, 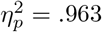) swing amplitudes. Treadmill symmetry conditions had no effect on arm swing asymmetry (*p* > .05) (Table 2).

**Table 1.**
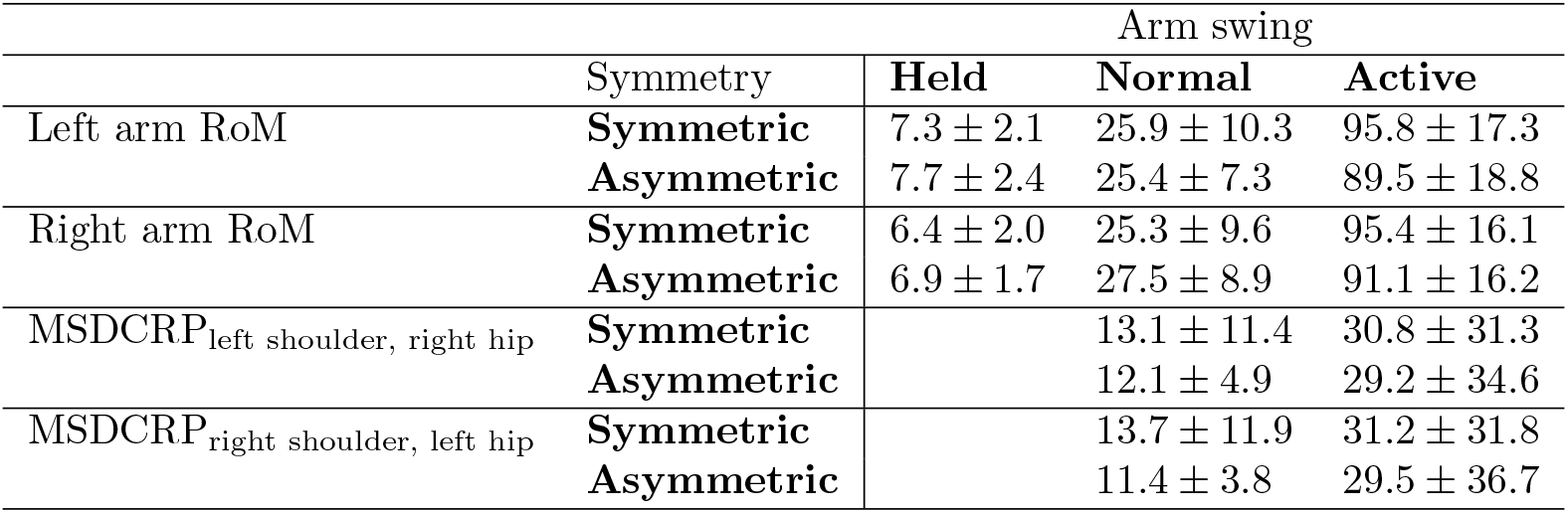
Summary of arm swing and contralateral limb coordination.

|  | Symmetry | Arm swing |  |  |
| --- | --- | --- | --- | --- |
|  |  | Held | Normal | Active |
| Left arm RoM | **Symmetric** | $7.3 \pm 2.1$ | $25.9 \pm 10.3$ | $95.8 \pm 17.3$ |
| | **Asymmetric** | $7.7 \pm 2.4$ | $25.4 \pm 7.3$ | $89.5 \pm 18.8$ |
| Right arm RoM | **Symmetric** | $6.4 \pm 2.0$ | $25.3 \pm 9.6$ | $95.4 \pm 16.1$ |
| | **Asymmetric** | $6.9 \pm 1.7$ | $27.5 \pm 8.9$ | $91.1 \pm 16.2$ |
| MSDCRP<sub>left shoulder, right hip</sub> | **Symmetric** | | $13.1 \pm 11.4$ | $30.8 \pm 31.3$ |
| | **Asymmetric** | | $12.1 \pm 4.9$ | $29.2 \pm 34.6$ |
| MSDCRP<sub>right shoulder, left hip</sub> | **Symmetric** | | $13.7 \pm 11.9$ | $31.2 \pm 31.8$ |
| | **Asymmetric** | | $11.4 \pm 3.8$ | $29.5 \pm 36.7$ |

**Table 2.** Summary of asymmetry measures and main effects.

|  | Symmetry | Arm swing |  |  | Main Effect |  |
| --- | --- | --- | --- | --- | --- | --- |
|  |  | Held | Normal | Active | Arm Swing | Symmetry |
| Arm swing asymmetry | **Symmetric** | $21.6 \pm 18.9$ | $22.1 \pm 14.1$ | $9.2 \pm 7.0$ | $F(2, 28) = 5.36, p = .011, \eta_p^2 = .277$ | n.s. |
| | **Asymmetric** | $23.0 \pm 12.8$ | $24.6 \pm 26.2$ | $10.9 \pm 7.9$ | | |
| Spatial step asymmetry* | **Symmetric** | $2.88 \pm 2.42$ | $3.73 \pm 2.31$ | $3.75 \pm 2.82$ | n.s. | $F(1, 14) = 90.37, p < .001, \eta_p^2 = .866$ |
| | **Asymmetric** | $16.42 \pm 6.18$ | $15.85 \pm 5.65$ | $13.85 \pm 5.69$ | | |
| Temporal step asymmetry† | **Symmetric** | $3.23 \pm 2.19$ | $3.23 \pm 2.02$ | $3.63 \pm 2.89$ | n.s. | n.s. |
| | **Asymmetric** | $1.94 \pm 1.24$ | $1.71 \pm 1.46$ | $3.41 \pm 2.12$ | | |

n.s. stands for not significant. \*Spatial step asymmetry had a significant interaction effect ( $F(2, 28) = 5.00, p = .014, \eta_p^2 = .263$ ). †Temporal step asymmetry approached significance for main effects of both arm swing ( $p = .064$ ) and symmetry ( $p = .054$ ).

MSDCRP between the left shoulder and right hip showed a significant effect of arm swing (*F* (1, 14) = 8.13, *p* = .013, 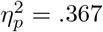); the MSDCRP between the right shoulder and left hip was also significantly affected by arm swing (*F* (1, 14) = 8.04, *p* = .013,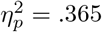). The MSDCRP of both contralateral limb pairs showed significant increases from normal to active swing (*p* = .013). Neither limb pair was affected by the gait symmetry conditions.

### Stability measures

See Table 3 for a summary of the significant main effects for the stability measures. Post-hoc tests showed that the active arm swing had significantly lower *λ*_*s*_ compared to both normal (*p* = .002) and held (*p* = .001) swing conditions. Asymmetric walking had significantly higher *λ_s_* than symmetric (*p* < .001).

**Table 3.** Summary of stability related measures and main effects.

|  | Symmetry | Arm swing |  |  | Main Effect |  |
| --- | --- | --- | --- | --- | --- | --- |
|  |  | Held | Normal | Active | Arm Swing | Symmetry |
| Local dynamic stability | Symmetric | 0.52 ± 0.03 | 0.51 ± 0.03 | 0.49 ± 0.04 | $F(2, 28) = 18.31, p < .001, \eta_p^2 = .567$ | $F(1, 14) = 37.55, p < .001, \eta_p^2 = .728$ |
|  | Asymmetric | 0.59 ± 0.06 | 0.56 ± 0.05 | 0.50 ± 0.03 |  |  |
| Step width | Symmetric | 0.19 ± 0.04 | 0.19 ± 0.04 | 0.20 ± 0.04 | n.s. | $F(1, 14) = 27.36, p < .001, \eta_p^2 = .661$ |
|  | Asymmetric | 0.21 ± 0.04 | 0.21 ± 0.04 | 0.21 ± 0.04 |  |  |
| Step width SD* | Symmetric | 0.017 ± 0.004 | 0.018 ± 0.005 | 0.022 ± 0.007 | $F(2, 28) = 17.16, p < .001, \eta_p^2 = .551$ | n.s. |
|  | Asymmetric | 0.017 ± 0.004 | 0.017 ± 0.004 | 0.020 ± 0.005 |  |  |
| Left step length | Symmetric | 0.63 ± 0.04 | 0.63 ± 0.03 | 0.70 ± 0.04 | $F(2, 28) = 40.69, p < .001, \eta_p^2 = .744$ | $F(1, 14) = 175.38, p < .001, \eta_p^2 = .926$ |
|  | Asymmetric | 0.55 ± 0.05 | 0.55 ± 0.03 | 0.61 ± 0.05 |  |  |
| Left step length SD | Symmetric | 0.018 ± 0.004 | 0.016 ± 0.003 | 0.024 ± 0.005 | $F(2, 26) = 4.95, p = .035, \eta_p^2 = .276$ | $F(1, 13) = 13.42, p = .003, \eta_p^2 = .508$ |
|  | Asymmetric | 0.023 ± 0.008 | 0.023 ± 0.008 | 0.027 ± 0.011 |  |  |
| Right step length | Symmetric | 0.64 ± 0.03 | 0.64 ± 0.03 | 0.70 ± 0.05 | $F(2, 28) = 18.05, p < .001, \eta_p^2 = .563$ | n.s. |
|  | Asymmetric | 0.64 ± 0.05 | 0.65 ± 0.04 | 0.70 ± 0.07 |  |  |
| Right step length SD | Symmetric | 0.019 ± 0.006 | 0.017 ± 0.003 | 0.028 ± 0.009 | $F(2, 28) = 7.70, p = .002, \eta_p^2 = .355$ | $F(1, 14) = 33.07, p < .001, \eta_p^2 = .703$ |
|  | Asymmetric | 0.027 ± 0.006 | 0.028 ± 0.015 | 0.033 ± 0.008 |  |  |

n.s. stands for not significant. \*A main effect of symmetry approached significance for step width SD ( $p = .064$ ).

Step width variability was increased during active swing compared to both held (*p* = .001) and normal (*p* = .004). Step width increased during split-belt walking conditions (*p* = .001). Active swing resulted in increased step length compared to held and normal, respectively, for the right (*p* = .002, *p* = .001), and left (*p* < .001) sides. Right step length increased during split-belt walking (*p* < .001). Active swing also showed increased right step length variability compared to held (*p* = .002) and normal (*p* = .003). Asymmetric walking resulted in increased step length variability compared to symmetric for the left (*p* = .003) and right (*p* < .001).

### Measures of asymmetry

See Table 2 for a summary of the significant main effects for the measures of asymmetry. Given the Arms×Symmetry interaction effect in spatial step asymmetry, post-hoc tests showed a significant decrease between the held and active swing conditions only while in the asymmetric walking condition (*p* = .025), and the split-belt walking was significantly increased from symmetric gait (*p* < .001). Arm swing asymmetry was significantly decreased in the active swing conditions as compared to held (*p* = .016) and normal (*p* = .035) swing. Temporal asymmetry, shown in Fig 1, contained one outlier which was removed and approached significance for main effect of both arm swing (*p* = .064) and symmetry (*p* = .054).

**Fig 1.**
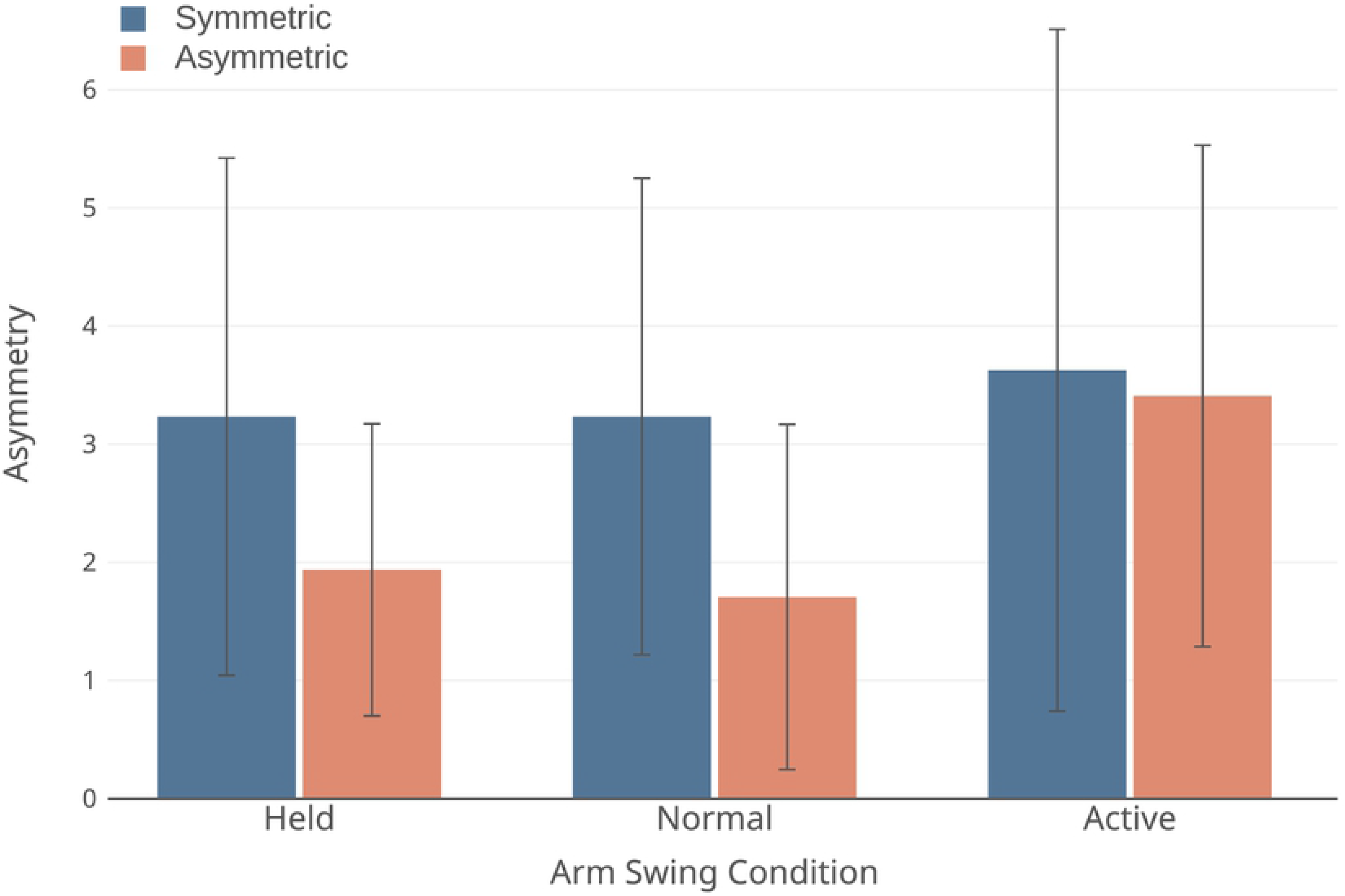
Temporal step asymmetry. Error bars are ±SD.

## Discussion

In this study, we examined the influence of changes in arm swing amplitude and lower limb asymmetry on gait stability using several stability metrics. When compared to both normal and held arm swings, active arm swing resulted in reduced contralateral limb coordination and increases in gait variability. However, the active swing also showed decreases in the maximal Lyapunov exponent. As for gait asymmetry, split-belt walking resulted in increased gait variability in spatiotemporal measures and maximal Lyapunov exponent. Altogether, while the difference in local dynamic stability between active and normal swings matches our hypothesis that active swing would increase local stability, it conflicts with both the increased step width and length variability and the decreased arm-leg coordination. In contrast, but in agreement with our hypothesis of asymmetric gait causing reductions in stability, both local dynamic stability and gait variability metrics show reduced stability during the split-belt walking conditions.

On the basis of these results, it is justified to revise the assumption that local dynamic stability and gait variability metrics are comparably related to stability, when defined as the likelihood of falling, and often referred to as global stability. While both types of measures have been linked to this definition of global stability in literature [18, 34, 35], the results here suggest a more nuanced application and interpretation of these metrics. The decreased MLE at the cost of increased gait variability demonstrates the importance of local trunk stability as a movement outcome-the trunk does account for a large proportion of the body mass. However, the addition of gait variability as a measure of stability complements the trunk local dynamic stability by providing a direct measure of the base of support which may signal decreases in global stability and increased risk of falling before the trunk signals reflect decreases in stability. Dingwell et al. (2000) found precisely this behaviour of a more locally stable trunk despite increased gait variability, and concluded that local dynamic stability and gait variability must be distinguished due to the fundamental differences of what they are classifying [36].

Therefore, the increased local dynamic stability of the trunk is in line with our hypothesis, increasing arm swing increases stability, if we clarify that stability here refers solely to the local stability of the trunk, which may change independently of the global stability. What remains unclear at this time is the cause of this increase in local trunk stability. We suggest two possible explanations, both of which may contribute to these independent changes in local trunk stability. Our initial hypothesis regarding increased arm swing, was primarily directed from a physics-based perspective, where increases in arm swing result in increases in angular momentum, thereby increasing the trunk’s resistance to change [37]. In this case, it is possible that the increased local stability from the greater angular momentum of the arms outweighs any decreases in arm swing coordination and increases in step variability. However, it is also possible that the more variable, or dynamic, nature of the base of support also contributed to a more locally stable trunk, but this would be difficult to confirm. A simulation study would be best suited to investigate this, however, it seems likely that replicating the behaviour seen here would require a model that actually demonstrates chaotic behaviour (i.e. the model can be mathematically represented as a dynamical system), as opposed to previously used passive dynamic walker models [35, 38].

With respect to the gait variability measures, we attribute the increase in gait variability between active swing and the other two conditions primarily to the decrease in inter-limb coordination. Decreased inter-limb coordination during active swing is also visible in a trend towards increased temporal step asymmetry when compared to normal swing. The decrease in coordination during active arm swing is likely due to a switch from automatic central pattern generator movements to conscious control (i.e. dual tasking) of arm swing. Active arm swing acting as a dual-task could also contribute to the increased gait variability. This behaviour would be in line with the results of McFadyen et al. (2009) who found changes in double support proportion and variation under the combined challenge of split-belt walking and a dual-task [39]. However, while the negative effect of dual-tasking on stability is well established in impaired populations [40, 41], this negative effect is not as well supported in young adults studied here [40, 42], and the design of this study is not appropriate to resolve this question.

Whatever the cause of the increased gait variability during active arm swing, the physical challenge of maintaining stability must increase with increasingly variable steps as increased gait variability represents a continuous minor perturbation to gait. It remains to be seen if more moderate increases in arm swing amplitude could succeed, or if prior training with this level of swing amplitude could mitigate any affect of dual-tasking or coordination issues. Further studies which include a continuum of arm swing amplitudes would be also insightful as to the relationship between swing amplitude and stability, and whether there may be a transition point beyond which increased swing amplitude decreases stability.

Regarding the effects of the split-belt walking, the induced asymmetric gait reduced stability as expected by increasing step length and width variability as well as reducing the trunk local stability. The spatial step length asymmetry was dramatically increased by the split-belt walking also as expected, while the temporal step asymmetry and arm swing symmetry were unchanged. The results of temporal symmetry in the lower limbs during split-belt walking agree with past research by Malone et al. (2012) which showed that both temporal and spatial asymmetry adapts to split-belt walking [43], and other studies showing that asymmetric gait requires more precise timing of gait patterns (i.e. temporal asymmetry should remain roughly the same or improve) [44, 45]. In our results, the trend towards improved temporal asymmetry in the held and normal swing conditions supports the previous results. Note that the spatial asymmetry calculated here differs from that used by Malone et al. (2012) and does not necessarily conflict with previous results regarding spatial adaptation [43]. The unchanged arm swing symmetry between symmetric and asymmetric gait is also in line with a previous study on upper and lower limb coordination during asymmetric gait which found that upper limb movements maintain symmetry and follow the rhythm of the fastest leg, even during increasingly asymmetric gait [45].

While it is possible that the decreased stability during the split-belt walking is due to the novelty, acting as a dual-task, previous research indicates that this is improbable [43, 44]. Indeed, Dietz et al. (1994) saw adaptations to inter-limb coordination in 10-20 strides during split-belt walking across multiple split-belt speed ratios [46], and Reisman et al. (2005) showed rapid changes to single limb characteristics, again across multiple speed ratios [44]. Despite the adaptability of multiple aspects of gait (including some measures of gait stability), some studies suggest that not all characteristics of gait stability can adapt to long-term split-belt walking [22, 23]. A thorough investigation on gait stability in adults familiar with split-belt walking could confirm the likely destabilizing effects of asymmetric gait.

In addition, healthy older adults have been shown to have reduced adaptation to asymmetric gait than healthy young adults [47]. Similarly, people with PD have demonstrated worse adaptability than age-matched controls [48]. While not the only population suffering from asymmetric gait, people with PD are one example of those who demonstrate a pathological gait with clear changes in arm swing amplitude and symmetry, and gait asymmetry when compared to age-matched controls, while also showing decreased stability (i.e. increased risk of falls) [49–51]. Our results suggest that deficits in stability may be linked to the decreases in gait symmetry rather than decreases in arm swing symmetry. Future work involving both induced asymmetric gait and asymmetric arm swing and their effects on stability could clarify the relative contributions of each towards gait stability.

## Conclusion

In conclusion, while active arm swing does improve local trunk stability, simultaneous increases in gait variability metrics suggest that: 1) local dynamic stability should not be treated as precisely analogous to standard gait variability variables due to the fundamental differences of what and how they characterize stability, and 2) active arm swing does not appear to be an appropriate intervention to decrease the likelihood of falling in healthy, young adults. As well, increases in lower, but not upper, limb asymmetry during split-belt walking indicates that the corresponding decreases in gait stability for asymmetric gaits are likely linked to the increased lower limb asymmetry–a finding relevant to pathological gaits with marked upper and lower limb asymmetries as well as decreased global stability.

